# PI3K regulates intraepithelial cell positioning through Rho GTP-ases in the developing neural tube

**DOI:** 10.1101/179325

**Authors:** Blanca Torroba, Antonio Herrera, Anghara Menendez, Sebastian Pons

**Author notes:** Correspondence author: Sebastian Pons, Instituto de Biología Molecular de Barcelona, (CSIC), Parc Científic de Barcelona, Baldiri Reixac 10-12, Barcelona 08028, Spain. Present Affiliation for Blanca Torroba: Department of Physiology, Anatomy and Genetics, University of Oxford, Le Gros Clark Building, S Parks Rd, Oxford OX1 3QX (UK).

## Abstract

During neural tube development, PI3K pathway promotes cell survival and provides the apical-basal navigation clues that define the final location of neurons in the epithelium.

**SUMMARY:** Phosphatidylinositol 3-kinases (PI3Ks) are signal transducers of many biological processes. Class 1A PI3Ks are hetero dimers formed by a regulatory and a catalytic subunit. We have used the developing chicken neural tube (NT) to study the roles played by PI3K during the process of cell proliferation and differentiation. Notably, we have observed that in addition to its well characterized anti apoptotic activity, PI3K also plays a crucial role in intra epithelial cell positioning, and unlike its role in survival that mainly depends on AKT, the activity in cell positioning is mediated by Rho GTPase family members. Additionally, we have observed that activating mutations of PI3K that are remarkably frequent in many human cancers, cause an unrestrained basal migration of the neuroepithelial cells that end up breaking through the basal membrane invading the surrounding mesenchymal tissue. The mechanism described in this work contribute not only to acquire a greater knowledge of the intraepithelial cell positioning process, but also give new clues on how activating mutations of PI3K contribute to cell invasion during the first stages of tumour dissemination.

## INTRODUCTION

The vertebrate Central Nervous System (CNS), derives from the neural plate, a region of the ectoderm on the dorsal surface of the embryo during gastrulation. The neuroepithelial (NEP) cells function as neural stem cells and differentiate, first, into the numerous types of nerve cells (neurons) and, later, into supportive cells (glia) present in the body (Kintner, 2002). NEP cells, like other epithelial cells, exhibit apical-basal polarity, with their apical plasma membrane lining the lumen of the neural tube while maintaining cell– cell adhesion and the integrity of the VZ, and their basal plasma membrane contacting the extracellular matrix (ECM), which demarcate the outer boundary of the neural tube (Gotz and Huttner, 2005). Specialized cell surface-associated ECMs, named basement membranes (BMs), underline epithelial (and neuroepithelial) cells at their basal surfaces (Roignot et al., 2013). The apical surfaces of individual NEP cells, known as apical junctional complexes (AJCs), are composed mainly of N-Cadherin-based adherens junctions (AJs), but not by tight junctions as observed in epithelial cells (Gotz and Huttner, 2005). Additionally, the basal membrane establishes focal adhesions based on Integrins with the ECM that can also trigger an outside-in signalling controlling cytoskeletal organization, force generation, differentiation and survival (Hood and Cheresh, 2002; Long et al., 2016). Establishment of apical-basal polarity involves cell-cell and cell-extracellular matrix interactions, polarity complexes and trafficking of membrane components to the apical or basolateral domain. Many protein families are involved in cell polarization, but also lipids like phosphoinositides that regulate cytoskeletal and membrane dynamics and trafficking (Gassama-Diagne and Payrastre, 2009). NEP cells are highly elongated and have a bipolar morphology with a ‘‘pearl-on-a-string’’ shape. In the early developing neural tube, the entire structure is mostly composed of a germinal neuroepithelium consisting of a rapidly dividing cell population with two remarkable interrelated features: pseudostratification and interkinetic nuclear migration (INM) (Taverna and Huttner, 2010). Pseudostratification refers to the fact that although all neuroepithelial cells extend from the luminal (apical) surface of the neuroepithelium to the basal lamina throughout their cell cycle, their nuclei are found at various positions along this apical-basal axis resulting in a multilayer appearance. And this is due to the INM, which refers to the fact that the nuclei of NEP cells migrate to different relative apical-basal positions during the cell cycle (Baye and Link, 2008). With the onset of neurogenesis, localization of neurons to their final destinations will form new basal layers of cells in addition to the germinal neuroepithelium (also called, ventricular zone VZ and later the ependyma). These basal layers become progressively thicker as more cells are added, assuming the names of intermediate layer (IL), where post-mitotic cells localize, the mantle zone (MZ), where cells differentiate into both neurons and glia and a cell-poor marginal zone containing mainly nerve fibres (Diez del Corral and Storey, 2001; Marthiens and ffrench-Constant, 2009). Phosphatidylinositols (PtdIns) can be phosphorylated in the 3, 4 and/or 5 position. PtdIns(4,5)P_2_ (PIP2) and PtdIns(3,4,5)P_3_ (PIP3) represent less than 1% of membrane phospholipids, yet they function in a remarkable number of crucial cellular processes (Czech, 2000; Katso et al., 2001; Vanhaesebroeck et al., 2001). PIP3 is the mediator of multiple downstream targets of the phosphoinositide-3-kinase (PI3K) pathway. PI3Ks are grouped into three classes (I – III), however class I PI3Ks have been the major focus of PI3K studies because they are the isoforms that are generally coupled to extracellular stimuli. Class IA PI3Ks are heterodimeric enzymes consisting of a catalytic subunit (p110) that associates with a regulatory subunit (Cantley, 2002). The vertebrate genome presents three genes (PIKCA, PIKCB, PIKCD) that code for three highly homologous Class IA catalytic subunits named as p110 (α, β, δ). All class IA catalytic subunits can associate equally with all of the class IA regulatory subunits. Two genes encode subunits of 85 kDa, termed p85α (PIK3R1) and p85β (PIK3R2), in addition The PIK3R1 gene also produces two major alternative transcripts encoding the smaller proteins p55α and p50α. A third gene (PIKR3) encodes p55γ, a protein with similar structure to p55α (Pons et al., 1995; Brachmann et al., 2005). Class IA PI3Ks are obligatory dimers in the cell, since no evidences for free p85 or p110 were found (Geering et al., 2007), accordingly overexpression of monomeric p110α had very low activity compared to its co-expression with a regulatory subunit (Yu et al., 1998). The Class IA regulatory subunits play three main roles on PI3K activity, stabilize the p110 subunit, inhibit p110 basal activity and trigger enzyme activation upon binding to stimulated membrane receptors (Hofmann and Jucker, 2012; Burke and Williams, 2015). We have used the developing chicken neural tube (NT) to study the roles played by PI3K during the process of cell proliferation and differentiation. Notably, we have observed that in addition to its well characterized anti apoptotic activity, PI3K also plays a crucial role in intra epithelial cell positioning, and unlike its role in survival that mainly depends on AKT, the activity in cell positioning is mediated by Rho GTPase family members. Additionally, we have observed that activating mutations of PI3K that are remarkably frequent in many human cancers, cause an unrestrained basal migration of the neuroepithelial cells that end up breaking through the basal membrane invading the surrounding mesenchymal tissue. The mechanism described in this work contribute not only to acquire a greater knowledge of the intraepithelial cell positioning process, but also give new clues on how activating mutations of PI3K contribute to cell invasion during the first stages of tumour dissemination.

## RESULTS

### Expression of class IA PI3Ks switches from the proliferative to the transition/mantle zone during neural tube development

All class IA PI3K subunits had previously been reported to be expressed at certain level in both the developing and mature mammalian nervous system through different approaches (Ito et al., 1995; Eickholt et al., 2007; Geering et al., 2007; Waite and Eickholt, 2010). We have assessed the expression of all class IA PI3K subunits (Fig 1A) at two different developmental stages of chicken spinal cord (Fig 1B) by *In situ* hybridization (IH) (Fig 1C,D). The chosen stages corresponded to 50-56 hours post fertilization (hpf; Hamburger-Hamilton [HH] stage 14-16), when most of mitosis are self-expanding proliferative divisions; and 96 hpf (HH stage 22), when neurogenesis has already began (Hamburger and Hamilton, 1992). Specific and strong signal was detected for all isoforms except for p110β, p110δ and p85β, which showed weak expression in the ventricular zone (VZ) at HH14-16 and almost no expression at later stages. In contrast, p110α, p85α, p55α, p50α and p55γ were highly expressed at HH14-16 in the whole neural tube excluding the most dorsal part, which corresponds to the Roof Plate (RP). This pattern changed with the onset of neurogenesis, restricting their expression at HH22 to the intermediate layer (IL), the mantle zone (MZ) and the RP. Notably p110α expression pattern was coincident with the pattern observed for the regulatory subunits, this is consistent with the fact that monomeric catalytic or regulatory PI3K subunits are not stable (Yu et al., 1998; Geering et al., 2007). Next we wondered whether mRNA expression was really reflecting the distribution of PI3K molecules. Thus, lacking antibodies that detected endogenous chicken PI3Ks, we transfected wild type bovine p110α (from now on called just p110α) to asses where was it stabilized (only dimers with regulatory and catalytic subunits are stable). Notably, transfected p110α was detected only at the MZ when expressed alone, but in the whole neural tube if p85α was co-expressed with it (Fig 2A). Similarly to the mRNA expression patterns, detection of transfected p110α changed from proliferative to differentiated neuron areas (HuC-D^+^ cells) during neural tube development (Fig 2 B-C). Note that the only non-differentiated group of cells that stabilized p110α at later stages (HH12 + 48 hours post electroporation (hpe)) was the V3 interneuron domain, a group of excitatory commissural interneurons right below the motor neuron domain with very high expression of p85α and p55α mRNA at HH22 stage (Fig 1D).

**Figure 1.**
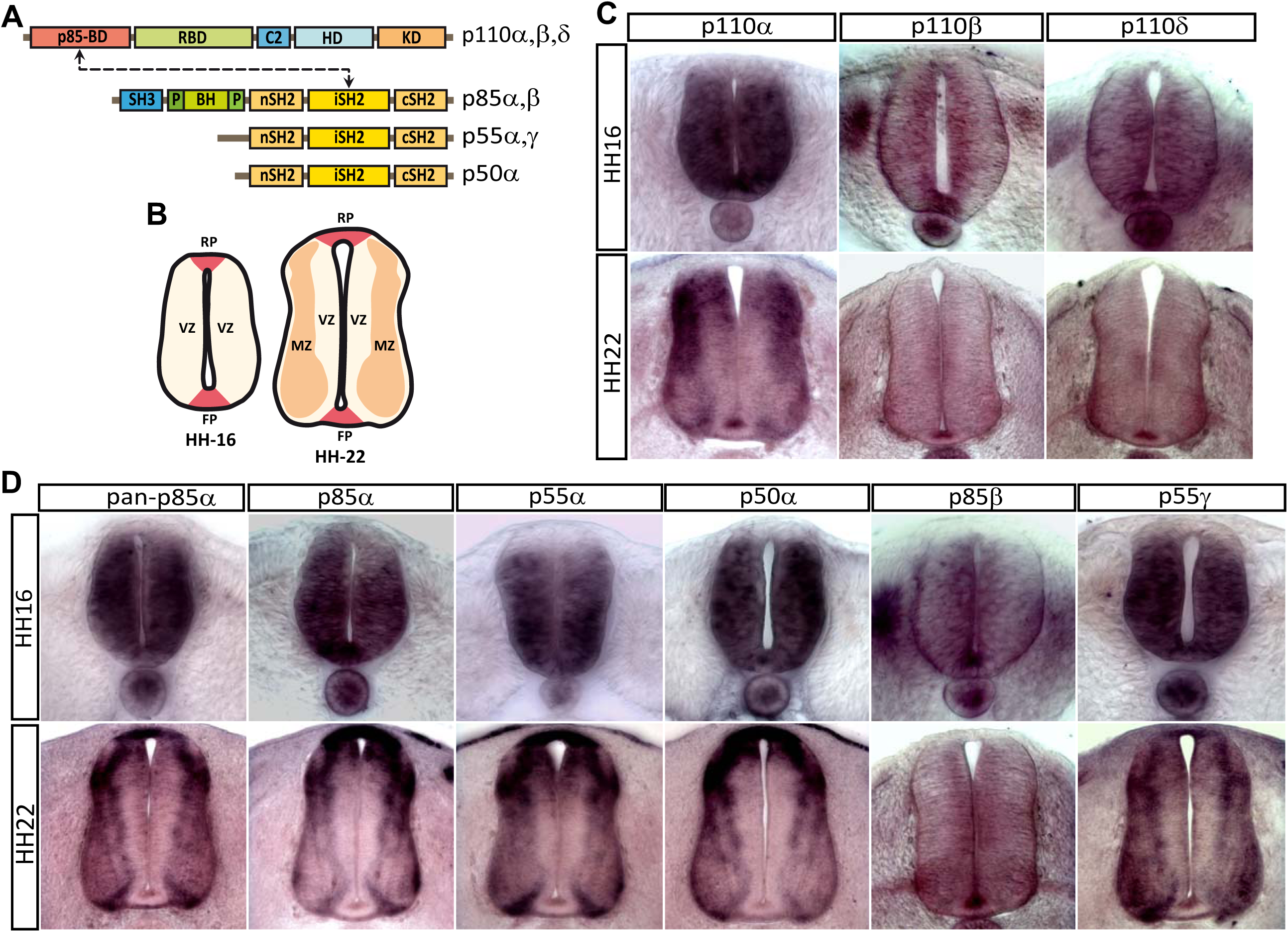
Class IA PI3K isoforms are differentially expressed at two different developmental stages of the chicken embryonic spinal cord. **(A)** Domain structure chart of catalytic and regulatory subunits of class IA PI3K **B)** Drawing of cell layer distribution of chicken spinal cord at HH12 and HH22. RP: roof plate, FP: floor plate, VZ: ventricular zone, MZ: mantle zone. **C,D)** In situ hybridization of class IA PI3K isoforms in chicken spinal cord transverse sections at HH stages 14-16 (50-56h) and 22 (96h)**. (C)** mRNA distribution of class IA PI3K catalytic subunits, *PIK3C1*, *PIK3C2* and *PIK3C3* corresponding to p110α, p110β and p110δ respectively. **(D)** mRNA distribution of Class IA PI3K regulatory subunits, *PIK3R1*,*PIK3R2* and *PIK3R3*. Note that *PIK3R1* gene generates p85α, p55α and p50α by differential splicing, whereas *PIK3R2* and *PIK3R3* give rise to p85β and p55γrespectively. Pan-p85α probe recognizes the three splice variants generated by the *PIK3R1* gene.

**Figure 2.**
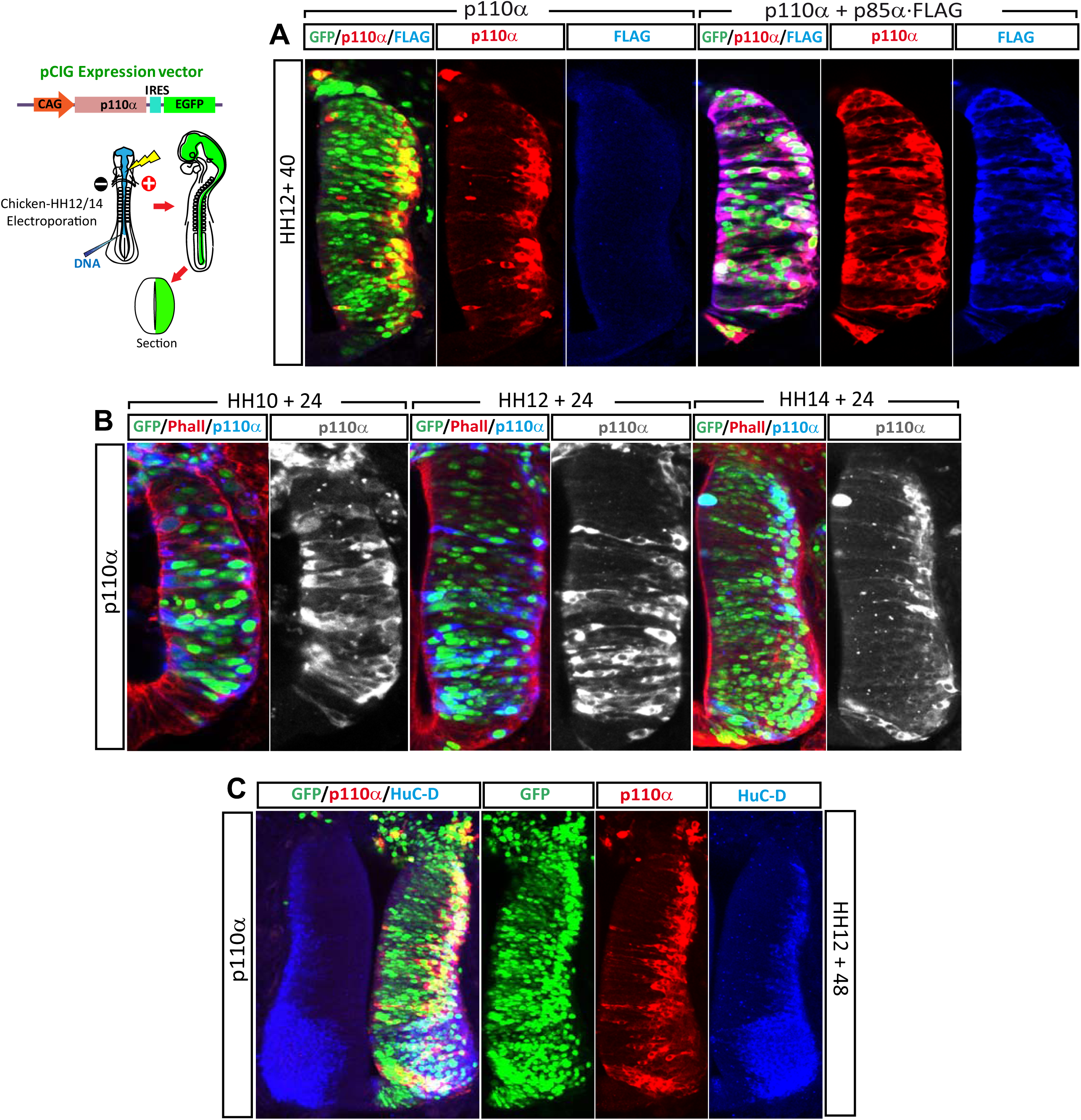
Transfection of bovine p110α reveals the distribution of Class IA PI3K regulatory subunits in the neural tube. **(A)** HH12 chicken neural tubes electroporated for 40h with p110α (wild-type bovine p110α cloned in the GFP expressing bicistronic vector pCIG) alone or in combination with FLAG·p85α. **(B)** HH10, HH12 or HH14 chicken neural tubes were electroporated with p110α for 24h, phalloidin was used to stain actin **(C)** HH12 chicken neural tubes were electroporated for 48h with p110α, HuC-D antibody was used to detect differentiated neurons. GFP staining denotes transfection. The antibody used to detect bovine p110α does not recognize the endogenous chicken form. Note that transfected p110α is detected only in the areas where it is stabilized, notably the same areas where the in situ hybridization predicted the presence of regulatory subunits. The bar graph represents the Mean ± SD of at least three independent electroporations, numbers in bars indicate the total counted cells.

### *PIK3CA* regulates survival of neuroblasts and neurons in the embryonic spinal cord

Neurotrophic factors as IGF-I or neurotrophins activate intracellular pathways including the PI3K pathway, which has been reported to be necessary and sufficient for survival of several neuronal types mainly through activation of the serine/threonine kinase Akt (Hetman et al., 1999; Brunet et al., 2001; Rodgers and Theibert, 2002). Moreover, Class IA PI3K have unique roles in mediating distinct neuronal functions such as protein synthesis, axonal outgrowth or synapse development and p110 dysregulation leads to neuronal dysfunction and apoptosis (Gross and Bassell, 2014). Therefore, we wondered whether p110α was needed for neural survival in this specific model. We knocked down p110α by transfecting HH-12 embryos with a mixture of three validated short hairpin inhibitory RNAs directed against p110α (shp110α) cloned in pSHIN, a GFP expressing bicistronic vector, and studied apoptosis 24, 48, 72 and 98 hpe. Notably, p110α knock down significantly increased apoptosis in the neuroblasts of the VZ 24 hpe (Fig 3A), and at longer electroporation times apoptotic cells were also observed at the MZ (Fig 3B), indicating that p110α had anti-apoptotic actions in neuroblasts and also in differentiated neurons. To confirm this, we studied the effect of p110α knock down on the number of Sox2^+^ (progenitors) and HuC/D^+^ (neurons) cells at two electroporation times. The knock down of p110α induced a reduction of Sox2^+^ cells of 42.04% at HH12+40 hpe and of 33.7% at HH12+72 hpe. Similarly HuC/D^+^ population fell by 56.66% at HH12+40 and 49.52% at HH12+72. At the two electroporation times studied the effect was greater on the neuron population; however this could be due to the cumulative effect.

**Figure 3.**
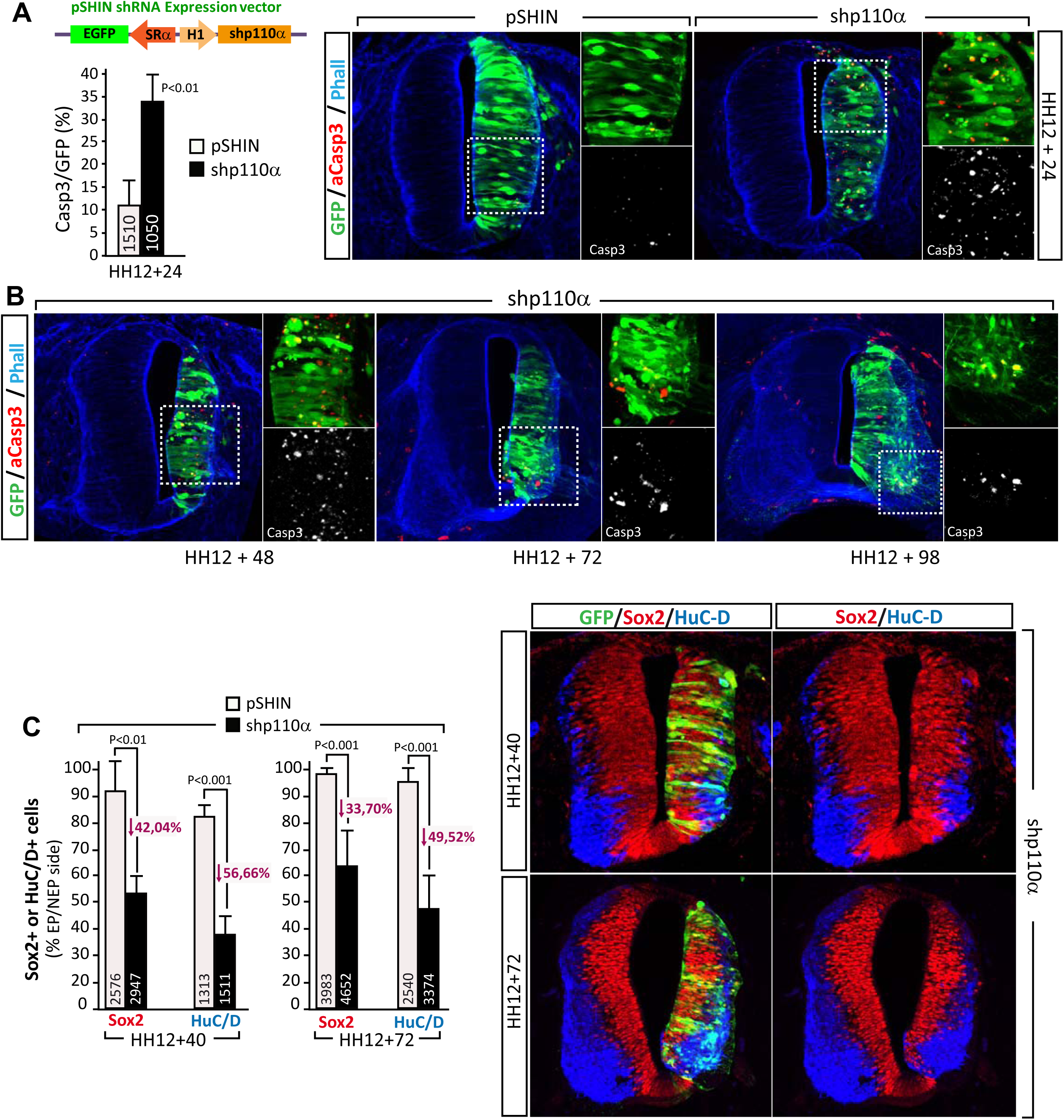
p110α prevents apoptosis in neuroblasts and neurons. **(A)** A short hairpin inhibitory-RNA directed against chicken *PIK3CA* (p110α) cloned in the GFP expressing vector pSHIN, was transfected in HH12 chicken neural tubes for 24h. Knock down of p110α significantly increased Caspase 3^+^ cells as compared to scrambled shRNA or empty vector transfected neural tubes. **(B)** Abundant Caspase 3^+^ cells were observed at the mantle zone at longer transfection times (48-72 and 98hpe). At 98hpe the motoneuron domain was particularly affected. The effect was cell autonomous, since only transfected cells (GFP^+^) were Caspase 3^+^. **(C)** Sox2^+^ (neuroblasts) and HuC-D^+^ (neurons) populations were evaluated at 40 and 72hpe of a p110α shRNA or scrambled shRNA, electroporated side (EP) was compared to non electroporated side (NEP), at both transfection times the reduction was greater in neurons than in neuroblasts, indicating a cumulative effect and confirming the effect of p110α knock down in both populations.

### *PIK3CA* regulates apical-basal positioning within the neuroepithelium

In addition to the pro-apoptotic effect, knock down of p110α also caused a miss location of cells in the neuroepithelium, where HuC/D^+^ cells (post mitotic cells differentiating into neurons) intermingled with Sox2^+^ (progenitors) instead of being accumulated at the MZ. This effect was cell autonomous because it was not observed in non-transfected cells (Fig 4A). We also observed that mature motoneurons (Isl-1^+^) failed to occupy the most lateral part of the motoneuron domain, differentiating instead at more apical positions (Fig 4A). Notably, a very similar result was obtained when p55α was knocked down (Fig 4B). As in p110α knock down, neuroblasts differentiated into neurons at more apical positions, leaving in many cases groups of Sox2^+^ progenitors trapped between the layer of differentiated neurons and the basal lamina (Fig 4B), and again, the motoneuron domain (Isl-1^+^ cells) did not reach the most basal positions. Knock down experiments using a shRNA construct targeting a common exon of *PIK3R1* (affects p85α, p55α and p50α) demonstrated similar effects than knocking down p110α or p55α (Fig 4C). Notably, the knock down of p110α for longer periods induced the differentiation of many neurons in the ventricle itself. The fact that these cells still projected normal axons heading towards the normal motoneuron exit point, indicated that a problem in intraepithelial cell positioning rather than a failure in survival or differentiation is what caused the abnormal layer distribution (Fig 4D).

**Figure 4.**
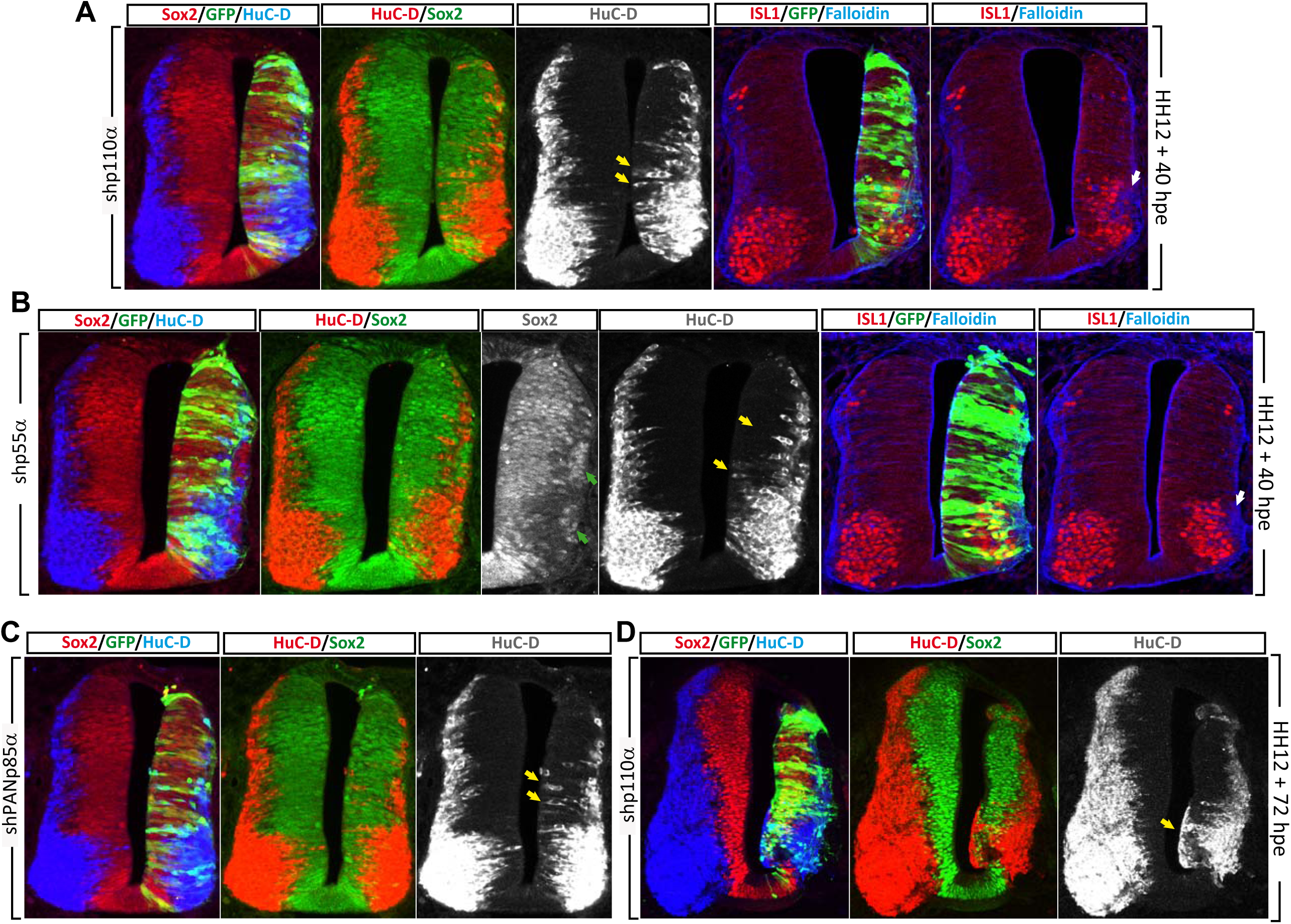
PI3K knock down impairs basal migration of differentiating neurons. **(A)** HH12 chicken neural tubes 40hpe with p110α shRNA show HuC-D^+^ differentiated neurons in the ventricular zone intermingled with Sox2^+^ neuroblasts (yellow arrows). Similarly staining with ISL1 motoneuron marker revealed that differentiated motoneurons did not reach the most basal positions (white arrow). **(B)** A similar experiment as in (A) but knocking down p55α, similarly an impairment of basal migration was observed, in this case groups of Sox2^+^ neuroblasts were frequently observed between the mantel zone and the basal membrane (green arrows). Motoneuron domain was also smaller and at more apical position compared to the non electroporated side. **(C)** A similar result was obtained with a shRNA targeting a common exon present in the three splice variants of *PIK3R1* (p85α, p55α, p50α). **(D)** A similar experiment as in (A) but at a longer electroporation time, failure in basal migration didn’t prevent differentiation, many neuros differentiated in contact with the lumen of the ventricle (yellow arrow). Pictures are representative of three independent electroporations. The effects were all cell autonomous since they were all restricted to electroporated cells (green).

Next, we performed gain-of-function experiments to obtain a deeper insight into the effects of PI3K on apical-basal cell positioning. No significant growth anomalies were observed in HH12 chicken neural tubes transfected separately with wild type (WT) p55α and p110α (not shown). However, mild but consistent growth aberrations occurred in the neural tubes electroporated with the combination of both subunits for 72 h (Fig 5A). Like so, groups of cells invaded the ventricle (blue arrowhead), and occupied the white matter located between the motoneuron domain and the basal membrane (red arrowhead), in consequence this space was diminished as compared to the non-transfected side. Notably, this same space was considerably enlarged when p110α was knocked down (Fig 5A). Next, we expressed p55α^W63R^, a constitutively active mutant recently described in our laboratory **(Supplementary Fig 1)**. Again, the effect of this mutant was mild when expressed alone (data not shown), however a strong phenotype was observed when p55α^W63R^ and p110α were co-transfected at an equimolar ratio. Although this mutation also induced the accumulation of cells in the ventricle, the most striking effect was a massive movement of cells towards the basal side (Fig 5B). Groups of cells that contained Sox2^+^ progenitors and HuC/D^+^ neurons headed towards the basal side moving initially as compact cell collections still restrained by the basal lamina (stained with laminin in Fig 5B) causing deformities of the neural tube (Fig 5C). However, in many cases the lamina was breached and the cells moved further (Fig 5B). These groups of cells maintained the N-Cadherin expression and did not mix with the surrounding mesenchymal cells; finally they fussed with the myotome which also expressed abundant N-Cadherin. Pictures in Fig 5B show two examples of this phenotype and a higher magnification capture of the bridges formed between the neural tube and the myotome. An identical result was obtained when p55γ^W65R^ or p85α^W333R^, the equivalent mutants in p55γ and p85α, were used **(Supplementary Fig 2)**. This massive cell movement occurred at all cranial-caudal levels studied and were not influenced by the position respect to the somites. Figure 5D shows a dorsal view of a whole mount preparation of a HH10 chicken embryo 40 hpe, where the space between the neural tube (NT) and the myotome (MT) is invaded by the cells electroporated with p55α^W63R^ plus p110α. We next wondered whether the same phenotype would be obtained by using p110α^H1047R^, a naturally occurring p110α mutation that is the most prevalent oncogenic mutation in breast cancer (http://cancer.sanger.ac.uk/cosmic) (Fig 5E). Interestingly when p110α^H1047R^ was combined with p55α or p85α the phenotype obtained was basically the same than with p55α^W63R^ plus p110α, consisting in cells invading the ventricle and also groups of cells breaching across the basal membrane, however when p110α^H1047R^ was used alone, the apical phenotype was absent (Fig 5E). This result is coherent with the fact that at HH12+40 h, p110α is stabilized only at the MZ (Fig 2) indicating that oncogenic forms of p110α that promote cell invasion still require the presence of equimolar concentrations of regulatory subunits to prevent their degradation.

**Figure 5.**
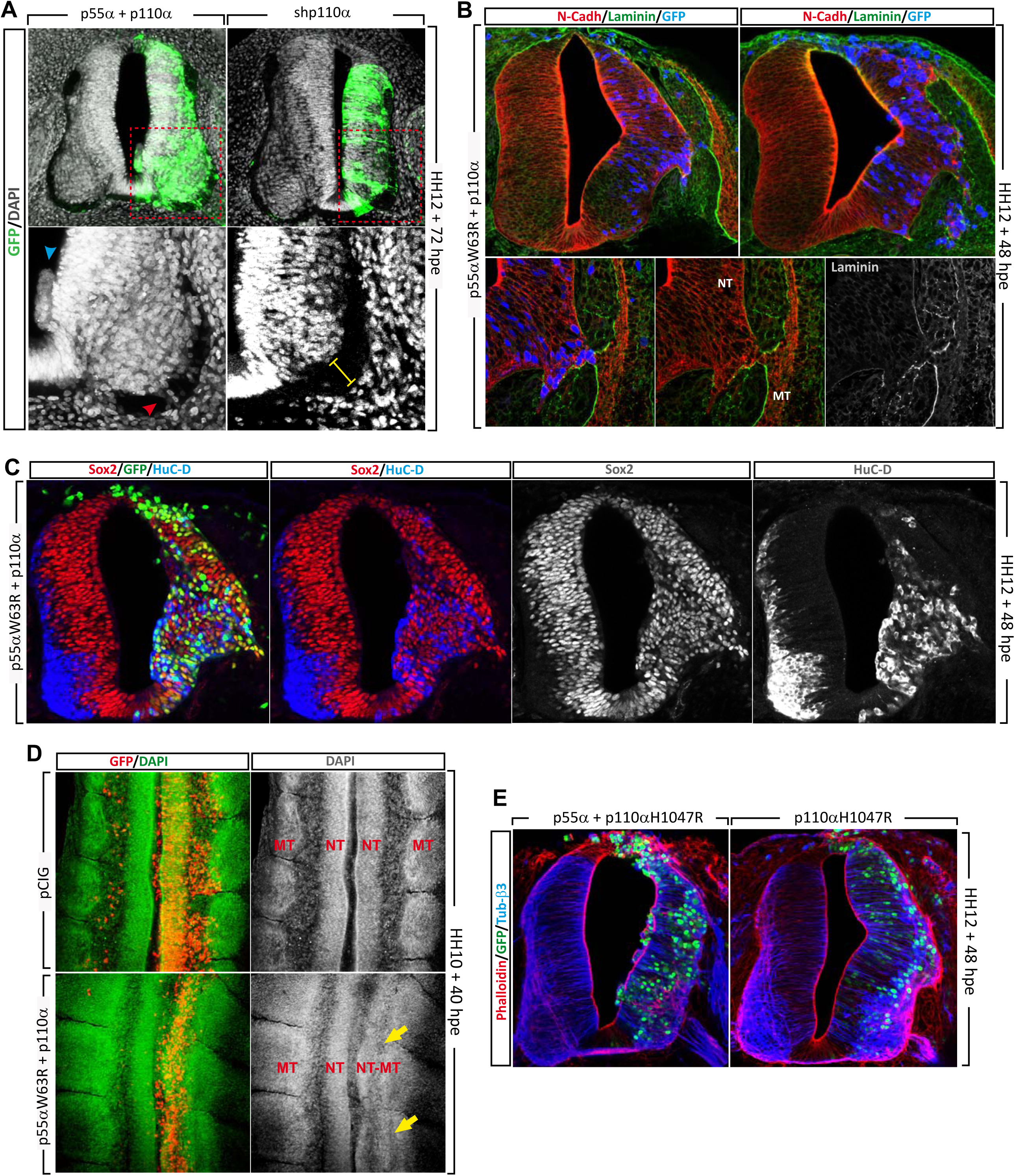
PI3K gain of function causes unrestraint basal migration. **(A)** HH12 chicken neural tubes transfected for 72h with WT p55α/p110α or a shRNA targeting p110α. GFP denote transfected cells, the area encircled by the red dotted line is enlarged in the lower panels. Cells invading the ventricle and the marginal zone are indicated by blue and red arrows respectively. The marginal zone is enlarged in the tubes where p110α is knocked down (yellow bar). **(B)** HH12 neural tubes were transfected for 48h with WT p110α plus p55α^W63R^ (active mutant), N-Cadherin labels the neural tube and the myotome, Laminin de basal membrane and GFP denotes the transfected cells. The lower panels show higher magnification images of transfected cells over migrating towards the basal side at the moment of breaching the basal membrane and fussing with the myotome. (NT: Neural Tube, MT: Myotome). **(C)** Neuroblasts and neurons 48hpe with WT p110α plus p55α^W63R^ were stained in HH12 neural tubes with Sox2 and HuC-D antibodies respectively. **(D)** Whole mount preparations of HH10 neural tubes 40hpe with p110α plus p55α^W63R^, or empty vector. Note that the neural tube (NT) is fussed to the myotome (MT) in the side expressing p55α^W63R^ (yellow arrows). **(E)** HH12 neural tubes were electroporated for 48h with WT p55α plus p110α^H1047R^ or p110α^H1047R^, differentiated neurons and actin filaments were stained with an antibody against Tubulin-β3 and phalloidin respectively. GFP denotes transfected cells.

### PI3Kα controls apical-basal polarity and basal migration through RhoGTPases

The serine/threonine kinase AKT binds PIP3 through its PH-domain. Recruitment of AKT to the plasma membrane is critical for neuronal polarity. Active forms of AKT are sufficient to induce multiple axon formation (Yoshimura et al., 2006). Although groups of cells invading the ventricle were frequently observed in neural tubes transfected with an active form of AKT, it did not reproduce the basal migration caused by active PI3K **(Supplementary Fig 3)**. Moreover, neither the apical invasions nor the basal exits induced by the expression of p55α^W63R^, were affected by the expression of a dominant negative form of AKT (Fig 6A,B). On the other hand, PI3K regulates Rho family GTPases (Cdc42, Rac1 and RhoA) through the PH domain-containing Rho GEFs (guanine nucleotide exchange factor) and GAPs (GTPase-activating proteins). Therefore, as Rho GTPases play key functions in cell migration and polarity, we wondered whether the apical and basal invasions induced by active PI3K could be mediated by this family of proteins. We expressed constitutively active forms of Cdc42, Rac1 and RhoA (Cdc42^G12V^, Rac1^Q61L^ and RhoA^Q63L^) in HH12 chicken embryos. Although phenotypes resembling PI3K gain of function could be observed in some slices transfected with Cdc42^G12V^ or Rac1^Q61L^ **(Supplementary Fig4)**, these active mutants demonstrated a very destructive activity on the neural tube and in most cases the overall structure was lost. Alternatively, we used dominant negative forms of Cdc42, Rac1 and RhoA (Cdc42^T17N^, Rac1^T19N^ and RhoA^T17N^) combined with active PI3K. Notably, we observed that both Cdc42^T17N^ and Rac1^T19N^ significantly reduced the number of slices showing basal exits, not affecting however the apical invasions of the ventricle. On the contrary, RhoA^T17N^ drastically reduced the ventricle invasions but had no effect on basal exits (Fig 6C,D). Moreover, combination of RhoA^T17N^ with Cdc42^T17N^ or Rac1^T19N^ had a complementary effect, reducing both apical and basal invasions (Fig 6E).

**Figure 6.**
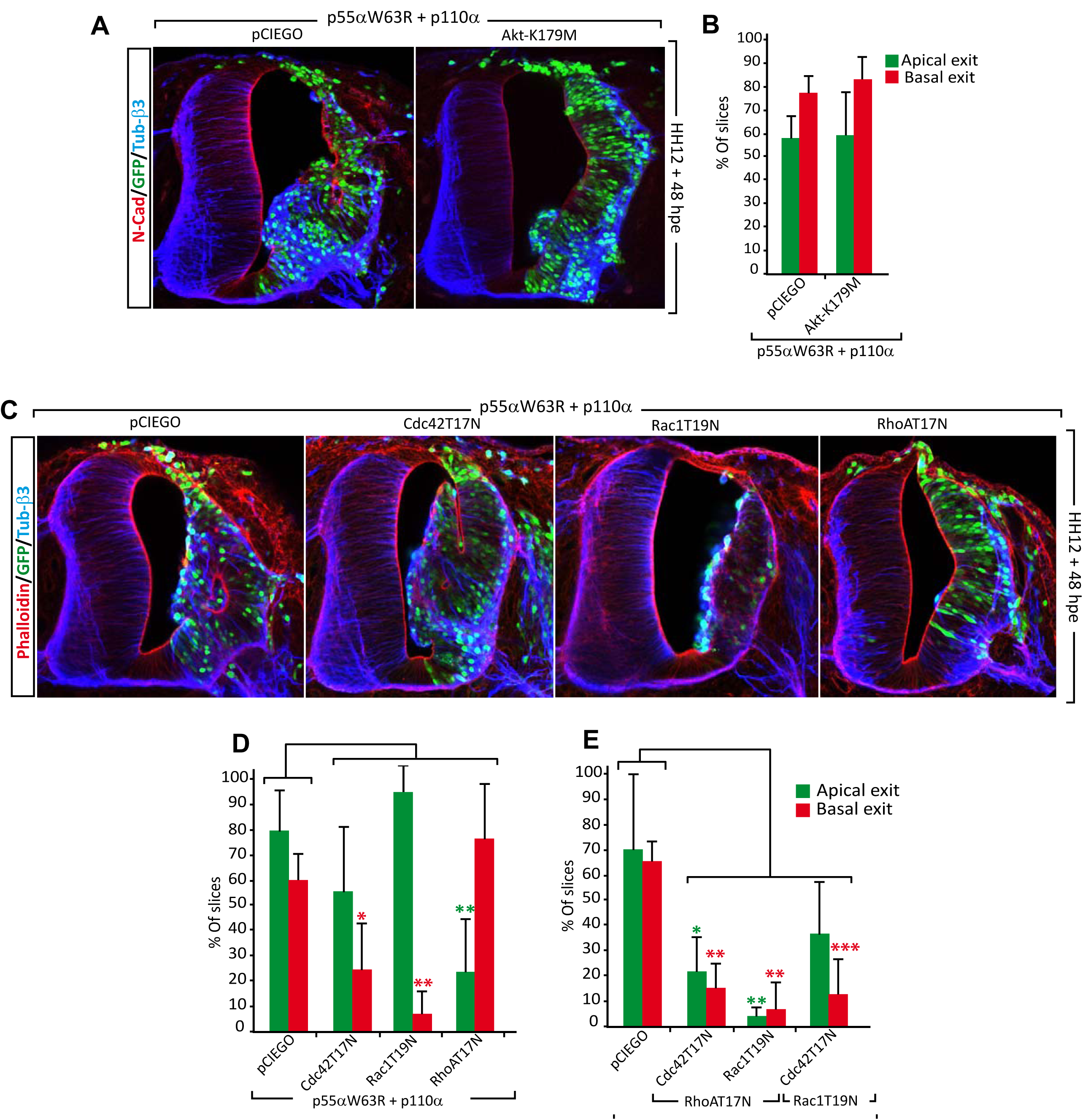
PI3K control cellular migration through Rho GTPases. **(A)** HH12 chicken neural tubes 48hpe with p110α, p55α^W63R^ and Akt^K179M^ (a DN mutant of Akt) or empty vector. The tubes were stained with anti N-Cadherin and Tubulin-β3 antibodies. GFP denotes transfected cells. **(B)** Akt^K179M^ expression did not modify the percentage of slices presenting apical or basal exits. **(C)** Experiment as in A where DN negative forms of Cdc42, Rac1 or RhoA were transfected together with p110α/p55α^W63R^. Differentiated neurons and actin filaments were stained with an antibody against Tubulin-β3 and phalloidin respectively. GFP denotes transfected cells. **(D)** Bar-graphs showing the percentage of slices presenting apical or basal exits for each transfection combination. Cdc42^T17N^ and Rac1^T19N^ significantly reduced basal exits whereas RhoA^T17N^ reduced apical exits. **(E)** Bar-graphs showing the effect of combined DN expression on the phenotype induced by p110α/ p55α^W63R^. The suppression activity of RhoA^T17N^ was complementary to Cdc42^T17N^ and Rac1^T19N^. The graphs represent the Mean ± SD of at least three independent elctroporations. *p<0.05, **p<0.01, ***p<0.001.

### Active PI3K induce uncontrolled apical and basal membrane protrusions

As mentioned before, transfection of different p110α/regulatory-subunit combinations caused mild phenotypes mainly consisting in cells invading the white matter adjacent to the motoneuron domain (Fig 5A). In addition, using membrane targeted GFP, we observed that some of the cells expressing p110α plus p55α produced basal processes that reached into the surrounding mesenchymal tissue at ectopic locations, but in this case none of the cell bodies invaded the adjacent mesenchyme (Fig 7A). In addition, we observed that endogenous PIP3 (studied with AKT-PH-GFP probe) concentrated in discrete spots at the apical and basal regions of the cell (Fig 7B). Interestingly, transfection of active PI3K (p55α^W63R^ plus p110α) increased the amount of PIP3 accumulated at the apical and basal poles, besides we observed that many of the cells accumulating PIP3 had oversized apical poles that protruded into the ventricle and invasive basal processes that breached the basal membrane and extended into the mesenchyme (Fig 7C-D). Notably, accumulation was especially intense in the segment of the basal process invading the mesenchyme (Fig 7D). To gather quantitative data, we divided each cell into ten segments, considering the first and the tenth as the apical and the basal segments respectively (Fig 7E), we measured the PIP3 level in the mentioned segments for each cell and divided it by the level measured in the lateral segments (2-9). Although active PI3K induced a significant accumulation of PIP3 only in the basal segment, a tendency to accumulate was also observed at the apical segment (Fig 7F). These results suggest causality between PIP3 accumulation and the unrestrained extension of the basal processes. Moreover, the invading filopodia could represent the mechanism through which the cells expressing active forms of PI3K initiate the invasion of the mesenchyme. Thus, without underestimating the contribution that the anti-apoptotic activity of PI3K may have on the initiation and progression of malignancies, we here propose that the uncontrolled cellular process induced by oncogenic forms of PI3K, could be crucial in the course of tumour invasion.

**Figure 7.**
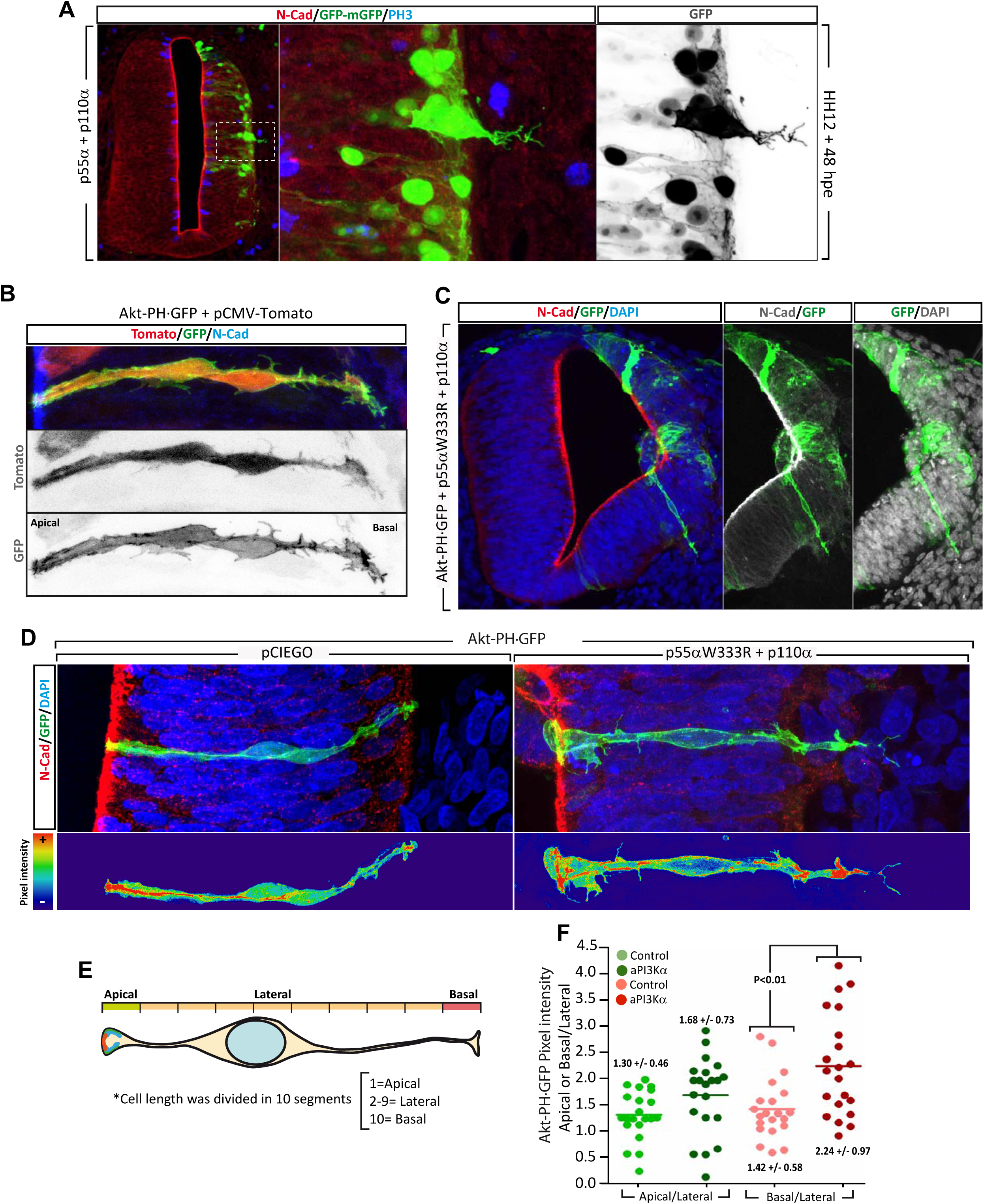
PI3K activity induces uncontrolled apical and basal membrane protrusions. **(A)** HH12 chicken neural tubes were transfected for 48h with p110α, p55α and membrane targeted GFP to visualize membranous processes in transfected cells. The tubes were stained with antibodies against N-Cadherin and P-histone H3 (labels M phase of mitosis). The encircled area is enlarged in the right panels. GFP channel is shown in grey scale to maximise contrast. The nuclear GFP corresponds to the pCIG vector in which p110α and p55α are cloned and denotes transfected cells. **(B)** HH12 chicken neural tubes were electroporated for 24h with Akt-PH·GFP probe to reveal PIP3 lipid distribution. Untargeted Red-Tomato was expressed as a control. Akt-PH·GFP accumulated at discrete spots along the neuroblast membrane including apical and basal locations as compared to the more diffuse distribution of Red-Tomato. Anti N-Cadherin was used to label the apical complex. **(C)** Akt-PH·GFP was transfected together with p110α/p55α^W63R^ or empty vector. Extra accumulation of PIP3 was observed in the invading processes induced by active PI3K. **(D)** Enlarged images of panel C and the corresponding controls transfected with empty vector. False color images are shown below indicating pixel intensity. **(E)** Scheme of quantification method for PIP3 accumulation using Akt-PH·GFP probe. The average pixel intensity was calculated at the apical, lateral and basal segments of each cell. Images were generated from HH12 neural tubes transfected for 24h with p110α/p55α^W63R^ or empty vector, 21 images coming from 3 independent transfections were used for each condition. **(F)** The scatter plot represents the apical and basal values divided by their corresponding lateral values. Active PI3K significantly increased PIP3 accumulation at the basal segment of the cell.

## DISCUSSION

The development of the neural tube is a complex process that requires coordinated morphogenetic forces together with localized extracellular signalling and controlled activation of genes involved in neural induction. In this work, we intend to decipher the role of Class IA PI3K proteins as signal transducers during early developmental stages of the spinal cord, which is anatomically the simplest and most conserved region of the vertebrate CNS and, therefore, it is a good model for studying neural polarity, cell division, neurogenesis or patterning among other processes.

### PI3K endure different biological activities along neural tube development

Although Class IA PI3Ks are known to be important for the development of the central nervous system, their expression pattern and molecular functions in the embryonic spinal cord has not been described before. In this work, we provide evidences of the participation of PI3Kα in cell survival, cell polarity and apical-basal positioning. More specifically, we show differential expression of catalytic and regulatory subunits before and after the onset of neurogenesis in the neural tube. Notably, at early stages the expression is abundant in the entire neuroepithelium, while at later stages Class IA PI3Ks become restricted to the neural population. We propose that this switch is associated to different functions along time; survival, mitosis and polarity at early stages and roles in migration and neural morphogenesis at later stages. Most cells in an organism are fated to die unless their survival is maintained by trophic factors produced either from neighbouring cells or provided by the bloodstream (Raff et al., 1993; Suzanne and Steller, 2013). PI3K has been reported to be a main survival pathway for many neuronal types; it mediates the activity of different growth factors signalling through RTKs (Receptor Tyrosine Kinase) such as Neurotrophins, Insulin, FGF or IGF-1 (Bartlett et al., 1991; Jungbluth et al., 1997). In agreement, we have demonstrated that both neuroblasts and neurons undergo apoptosis upon PI3K withdrawal in the embryonic chicken neural tube, indicating that although the anti-apoptotic signals acting at these two development stages are most likely different, PI3K remains as an essential anti-apoptotic transducer.

Previous articles linked PIP3 production with the establishment of apical-basal polarity in epithelial cells where PIP3 accumulation contributed to basolateral membrane determination. Segregation of PIP2 and PIP3 in the plasma membrane seems to be crucial to define apical and basolateral membrane, in MDCK (Madin-Darby canine kidney) cysts PIP2 becomes enriched at the apical membrane domain delimiting the lumen during cyst formation by recruitment of apical proteins such as annexin 2 or Cdc42, which activate the Par complex. In contrast, PIP3 is restricted to and specifies the basolateral surfaces of these cells through localization/interaction of PI3K with basal protein complexes like Dlg or after contacting with laminin surfaces (Martin-Belmonte and Mostov, 2008; Gassama-Diagne and Payrastre, 2009; Roignot et al., 2013). Here, we showed that alterations in PI3K activity severely disrupted neural tube architecture. Active-PI3K overexpression at HH12 disrupted the apical-basal polarity of the NEP as well as the distribution of apical complex proteins (Ncadherin, aPKC, ZO1) already after 17hpe, later on it induced cell-autonomous neuroepithelial malformations characterized by the presence of cell masses inside the ventricle and miss localization of mitotic cells at any position in the NEP, as previously reported after ectopic accumulation of apical markers like Par3 or aPKC (Afonso and Henrique, 2006; Herrera et al., 2014). Although it is conceptually difficult to separate the effect on intraepithelial cell positioning from the aberrant cell polarity, we believe that these are two mechanisms independently regulated by PI3K. Consequently, although PI3K knock down and the expression of active PI3K both induce cell depolarization, the effect on intraepithelial cell positioning is just the opposite, where knock down induces the cells to differentiate at half way to the marginal zone and overexpression provokes an excessive basal migration that ends up with neuroblasts breaking through the basal membrane and invading the surrounding mesenchyme. Thus, the concentration of PIP2 and PIP3 at the apical and basal poles of the cell needs to be finely tuned to achieve the correct location.

### Local control of PI3K activity restrains neuroblasts into the neuroepithelium

The next question would be how PIP3-dependent basal migration and axonal growth are endogenously regulated to avoid over-migration. A good candidate to delimit the CNS border would be the transmembrane guidance cue Semaphorin6A. Previous work has related Semaphorin expression with control of axonal growth in DRG neurons and spinal MNs (Mauti et al., 2007). Semaphorins are a large family of secreted or membrane-bound proteins shown to regulate axonal pathfinding during development of the nervous system through growth cone collapse, axon repulsion, or growth cone turning by regulating Rho GTPases and suppressing PI3K signalling pathway (Nakamura et al., 2000; Menager et al., 2004). During chick spinal cord development, Sema6A has been found at the transition zone between the peripheral and the central nervous system avoiding emigration of motoneurons (Mauti et al., 2007). Therefore, we hypothesize that under normal conditions PI3Kα activity is controlled by the Semaphorins coating the neural tube, avoiding neural somas to stream out throughout the DREZs (Dorsal Root Entry Zone) or MEPs (Motoneuron Exit Poins). However, over activation of PI3Kα might allow the cells to surpass the physical (basement membrane) and signaling based (Sema6A) barriers.

### PI3K control cellular migration through Rho GTPases

PI3K has been previously related to migration and invasiveness in different systems (Cain and Ridley, 2009; Rorth, 2011) including neurons where it has been implicated in Reelin-dependent migration (Kubasak et al., 2004; Valiente and Marin, 2010; Lee and D’Arcangelo, 2016). Our results suggest that PI3K mediates the activity of a basally-directing signal participating in the embryonic spinal cord lamination during neurogenesis. On the other hand, we have shown that PI3K induce cytoskeletal modifications involving F-actin and the expression of neuron specific beta-III tubulin. Several pathways downstream PI3K have been related to migration and/or actin based protrusions, among those, Rho GTPase family members play a central role in cell polarization and migration of several cell types including neurons (Iden and Collard, 2008; Azzarelli et al., 2014).

Notably, the expression of dominant-negative forms of Cdc42 and Rac1 but not of Rho A, significantly reduced the basally directed over-migration caused by active forms of PI3K. Interestingly, both Rac1 and Cdc42 dominant negative constructs also impeded the membrane protrusions into the mesenchyme and the basal lamina breaches associated to active PI3K. On the other hand, neither Rac1 nor Cdc42 dominant negatives molecules prevented the invasion of the ventricle. Complementarily, RhoA dominant negative molecules prevented ventricle occupation but had no effect on basal migration. These results suggest specific roles of RhoA versus Cdc42/Rac1 at apical and basal poles of the neuroblasts in mediating PI3K signaling. Interestingly, these topological differences resemble the disposition and activity of these three GTPases during cell migration, where generation of front-rear polarity is achieved by the local activation of Cdc42 and Rac1 at the front of the cell and RhoA at the back, resulting in rapid and dynamic assembly of actin filaments at the front and the assembly and activation of contractile actomyosin networks at the back (Nelson, 2009). Nevertheless, more studies are needed to understand how these cytoskeletal regulators are adapted to jump from apical-basal polarity to front-rear polarity regulation during migration, since Cdc42 is needed apically for AJC formation (Chen et al., 1999; Cappello et al., 2006) and subsequently in basal axonal growth cones (Grunwald and Klein, 2002; Meyer and Feldman, 2002). In this work, we offer evidences that link basal PIP3 accumulation and a requirement of Cdc42/Rac1 for basal neural migration during differentiation in the embryonic spinal cord.

### Role of PI3K in cancer invasion

*PIK3C1* and *PIK3R1,* coding for p110α and p85α respectively, are practically the only PI3K subunits mutated in cancer. Most of the mutations are concentrated at hot spots in the helical or catalytic domains of p110α and in the NSH2 or in the inter-SH2 domains of P85α, rendering in all cases constitutively active dimers of p110α and p85α. Mutation of *PIK3C1* is considerably more frequent than *PIK3R1*, in fact, *PIK3C1* is the most frequently mutated gene in breast cancer, being 110α^H1047R^ the predominant mutation. Notably, the incidence of this mutation is higher in invasive ductal breast carcinoma (IDC) than in ductal breast carcinoma in situ (DCIS) (http://cancer.sanger.ac.uk/cosmic), suggesting a role of this mutation in cancer progression and invasiveness. AKT is one of the main targets of PI3K pathway, mediating the anti-apoptotic activity of many growth factors. This fact has greatly influenced the current view regarding the relevance of each cellular process triggered by oncogenic PI3K on tumour initiation and progression. Consequently, anti-apoptosis may well be the main cellular process through which increased PI3K activity is believed to drive oncogenic growth. To the development of this idea also contributed the fact that many advanced carcinomas present inactivating mutations of PTEN, the main negative regulator of PI3K pathway, which activity is clearly pro-apoptotic. However, just as mutation of *PIK3CA* is particularly frequent in breast cancer, PTEN mutations are rare (http://cancer.sanger.ac.uk/cosmic), indicating that in addition to the anti-apoptotic effect, other cellular processes triggered by active PI3K mast be relevant to cancer progression. Developing neuroepithelium shares many common characteristics with breast ductal epithelium and with most of epitheliums in the body. In particular, breast ducts undergoes periodic cycles of growth and involution, where basal cells proliferate and differentiate into milk producing cells in each menstrual cycle, therefore acquisition of activating mutations of PI3K would not only prevent apoptosis after the hormone burst is over, but would also prone the mutant cells to migrate beyond the basal membrane limits contributing to the progression from *insitu* to invasive carcinoma. In brief, we believe that the results generated in this work using the neuroepithelium as a model may well be extrapolated to many other epitheliums susceptible of malignant transformation.

## MATERIALS AND METHODS

### Reagents

Antibodies, reagents and DNAs used in this work can be found in Supplementary Methods.

### In situ hybridization

Embryos were fixed 4hrs or overnight at 4°C in 4% PFA in PBS, rinsed and processed for whole mount RNA in situ hybridization following standard procedures using probes for chicken PI3K isoforms p110α, p110β, p110δ, pan85α (recognizes the three isoforms from the gen *Pik3r1*: p85α, p55α, p50α), p85α, p55α, p50α, p85β and p55γ. They were designed from mRNA differential sequences for each gene obtained from the *Gallus gallus* genome (NCBI) and they were around 250bp. Hybridization was revealed by alkaline phosphatase-coupled anti-digoxigenin Fab fragments (Boehringer Mannheim). Hybridized embryos were rinsed in PBT (1%), post-fixed in 4% PFA, vibratome sectioned and photographed on a Leica DMR Microscope. The complete sequence of all the probes used can be found in Supplementary Methods.

### **Chick embryo *in ovo* electroporation**

Eggs from White-Leghorn chickens were incubated at 38.5°C in an atmosphere of 70% humidity. Embryos were staged according to Hamburger and Hamilton (HH). Chick embryos were electroporated with affinity purified plasmid DNA (1-2 μg/μl) in H_2_O with Fast Green (0.5 μg/μl). Briefly, plasmid DNA was injected into the lumen of HH stage 10-12 neural tubes, electrodes were placed either side of the neural tube and electroporation carried out using and Intracel Dual Pulse (TSS10) electroporator delivering five 50 ms square pulses of 20 V for HH-10 and 25 V for HH-12. Transfected embryos were allowed to develop to the specific stages, then dissected, fixed and processed for immunohistochemistry or in situ hybridization. For bromodeoxyuridine (BrdU) labeling, 5 μg/μl BrdU was injected into the neural tubes 4 hours prior fixation.

### Immunohistochemistry and BrdU incorporation

Embryos were fixed for 4 hours (up to HH-20) or overnight (bigger than HH-20) at 4°C in 4% paraformaldehyde and immunostaining was performed on vibratome (40 μm) sections following standard procedures. For BrdU detection, sections were incubated in 2N HCl for 30 minutes and then rinsed with 0.1M Na_2_B_4_O_7_ [pH 8.5]. After washing in PBS-0.1% Triton, the sections were incubated with the appropriate primary antibodies and developed with Alexa or Cyanine conjugated secondary antibodies. After staining, the sections were mounted and examined on a Leica SP5 confocal microscope.

### Immunoblotting

For immunoblotting experiments, both sides of the neural tube were electroporated sequentially. GFP^+^ areas of electroporated neural tubes were dissected under a fluorescence binocular microscope, dissolved in 1X SDS Laemmli sample buffer (5 embryos per condition) and sonicated. The insoluble material was removed by centrifugation and the samples were resolved by SDS-PAGE and transferred to nitrocellulose membranes. The membranes were blocked with 8% non-fat dry milk in TTBS (150 mM NaCl, 0.05% Tween-20 and 20 mM Tris-HCl [pH 7.4]) and incubated with primary antibodies. The membranes were developed with either A/G-coupled peroxidas using the ECL system, or with Alexa labelled secondary antibodies which were detected with the Odyssey Infrared Imaging System from LI-COR.

### RT-qPCR

Both sides of the neural tube were electroporated with shRNA or the control vector (pSHIN) at HH12. GFP^+^ areas were dissected dissected 24hrs later. RNA was extracted and purified using High Pure RNA Isolation Kit (Roche) and total RNA was analyzed using NanoDrop (NanoDrop ND-1000 Spectophotometer, ThermoScientific). Similar amounts of RNA were retro-transcribed using the High Capacity cDNA Reverse Transcription Kit (Applied Biosystems) and the qPCR performed in a LightCycler480 System (Roche) using SYBR Green (Bio-Rad). Oligonucleotides specific for chick glyceraldehyde 3-phosphate dehydrogenase (GAPDH) were used for normalization.

### Statistical analysis

For the statistical analysis it was used the GraphPad Prism software (version 4.0). Quantitative data are presented as the mean ± SD from at least three independent experiments. One-way ANOVA followed by the Tukey’s test or Student’s t-test were used to determine significance using 95% of confidence interval.

## ADDITIONAL INFORMATION

### Acknowledgements

The authors wish to thank all the researchers mentioned in Supplementary Methods for providing DNAs. Monoclonal antibodies were obtained from the Developmental Studies Hybridoma Bank, developed under the auspices of the NICHD and maintained by The University of Iowa, Department of Biological Sciences, Iowa City, IA 52242.

